# Cooperative Amyloid Fibre Binding and Disassembly by the Hsp70 disaggregase

**DOI:** 10.1101/2021.08.29.458036

**Authors:** J. G. Beton, J Monistrol, A Wentink, EC Johnston, AJ Roberts, B Bukau, BW Hoogenboom, HR Saibil

## Abstract

Although amyloid fibres are highly stable protein aggregates, a specific combination of human Hsp70 system chaperones can disassemble them, including fibres formed of α-synuclein, huntingtin or Tau. Disaggregation requires the ATPase activity of the constitutively expressed Hsp70, Hsc70, together with the J domain protein DNAJB1 and the nucleotide exchange factor Apg2. Recruitment and clustering of Hsc70 on the fibrils appear to be necessary for disassembly.

Here we use atomic force microscopy (AFM) to show that segments of *in vitro* assembled α-synuclein fibrils are first coated with chaperones and then undergo bursts of rapid, unidirectional disassembly. Cryo-electron tomography reveals fibrils with regions of densely bound chaperones extending from the fibre surface, preferentially at one end of the fibre. Sub-stoichiometric amounts of Apg2 relative to Hsc70 dramatically increase recruitment of Hsc70 to the fibres, creating localised active zones that then undergo rapid disassembly at a rate of ∼4 subunits per second.

## Introduction

Neurodegenerative diseases are characterised by progressive aggregation of specific proteins, such as α-synuclein (αSyn) in Parkinson’s disease, followed by neuronal cell death. Hereditary neurodegenerative diseases are caused by mutations that increase the cellular concentration or aggregation propensity of such proteins, suggesting a causal relationship between pathological aggregation and cell death. These pathological aggregates can form amorphous assemblies (Shahmoradian et al., 2019) and highly ordered cross-β amyloid fibres (Fitzpatrick et al., 2017; Schweighauser et al., 2020). The identity of the toxic species remains unknown, with evidence for toxicity of smaller, disordered oligomers, thought to be precursors to amyloid fibres (Baglioni et al., 2006; Winner et al., 2011), and for toxicity of amyloid fibres (Peelaerts et al., 2015), which may accelerate disease by nucleating further oligomerisation (Guo et al., 2013; Padrick and Miranker, 2002). Given their strong association with disease, it seems likely that clearance of amyloid fibres is important to prevent disease onset (Kuo et al., 2013; Nagy et al., 2016).

Disaggregation in metazoans is mediated by Hsp70, which uses its ATP-driven conformational cycle to dissolve aggregates (Gao et al., 2015; Mayer and Bukau, 2005; Nillegoda et al., 2015). Control of this Hsp70 ATPase cycle is provided by cognate J-domain protein (JDP) and nucleotide exchange factor (NEF) co-chaperones, which functionally tailor Hsp70s by controlling substrate selection and ATPase cycle progression. Indeed, a specific combination of the JDP DNAJB1, Hsc70 (the constitutively expressed human cytosolic Hsp70) and Apg2 (Hsp110 type NEF), is sufficient to disaggregate Tau, αSyn and huntingtin exon 1 amyloid fibres (Gao et al., 2015; Nachman et al., 2020; Scior et al., 2018). These amyloids are thought to be disassembled by Hsc70/DNAJB1/Apg2 derived entropic pulling, caused by the entropic penalty that arises from steric clashes between chaperones bound to fibres at high density (Wentink et al., 2020).

DNAJB1 recruits Hsp70 to fibres and cannot be substituted by other JDPs in amyloid disaggregation (Gao et al., 2015; Wentink et al., 2020). This recruitment occurs through a 2-step mechanism initiated when the Hsp70 C-terminal EEVD motif binds to DNAJB1. This binding releases the auto-inhibition of DNAJB1’s J-domain, which is then free to bind the Hsp70 ATPase domain, stimulating ATP hydrolysis (Faust et al., 2020). This 2-step binding process is necessary for DNAJB1-facilitated clustering of Hsp70 on αSyn amyloid fibres, which is in turn necessary for disaggregation (Faust et al., 2020; Wentink et al., 2020). The force generated by the Hsc70/DNAJB1/Apg2 disaggregase is sufficient to fragment or depolymerise αSyn amyloid fibres, resulting in disaggregation (Gao et al., 2015). However, the dynamics of disaggregation are unknown. It remains unclear how the chaperones are organised at the fibre surface and how the amyloid fibres are remodelled during disaggregation.

Here, we explore how the actions of the Hsc70/DNAJB1/Apg2 disaggregase lead to the disassembly of amyloid fibres, using αSyn fibres as a model substrate. We used AFM to visualise the disassembly process in real time. The resulting videos show how disaggregation proceeds by combined fragmentation and depolymerisation of individual fibres and reveal regions of long-range chaperone clustering to be a necessary precursor for processive disassembly. Binding and activity assays, fluorescence microscopy and cryo-electron microscopy (cryo-EM) show that Apg2 recruits dense clusters of Hsc70 to the fibres, preferentially at one end, priming rapid bursts of disaggregation.

## Results

### Time-resolved AFM of amyloid fibre disaggregation

To image the disaggregation process by AFM, we developed a system to adsorb αSyn fibres on a mica surface while avoiding excessive adsorption of the Hsc70/DNAJB1/Apg2 chaperones. We achieved this by incubating mica surfaces first with αSyn fibres and then with a PLL-PEG copolymer. This PLL-PEG has been shown to reduce non-specific protein binding to mica surfaces in AFM experiments (Akpinar et al., 2019): the PLL component binds the negatively charged mica surface whilst the PEG component acts as a steric barrier to protein binding. Using this preparation, we could visualise αSyn fibres at the sample surface and greatly reduced non-specific adsorption of subsequently added chaperones onto the PEG-PLL passivated substrate. This enabled us to monitor the specific effect of added Hsc70/DNAJB1/Apg2 and ATP on αSyn fibres, including their resulting disaggregation. This disaggregation was visualised in 8 video series in 8 independent AFM experiments (Figure 1A, B), with a temporal resolution of 1-3 minutes per frame. This temporal resolution was suitable for stably imaging the surfaces over the longer time scales required to observe disaggregation events, yet was still fast enough to distinguish different steps in the disaggregation process.

**Figure 1.**
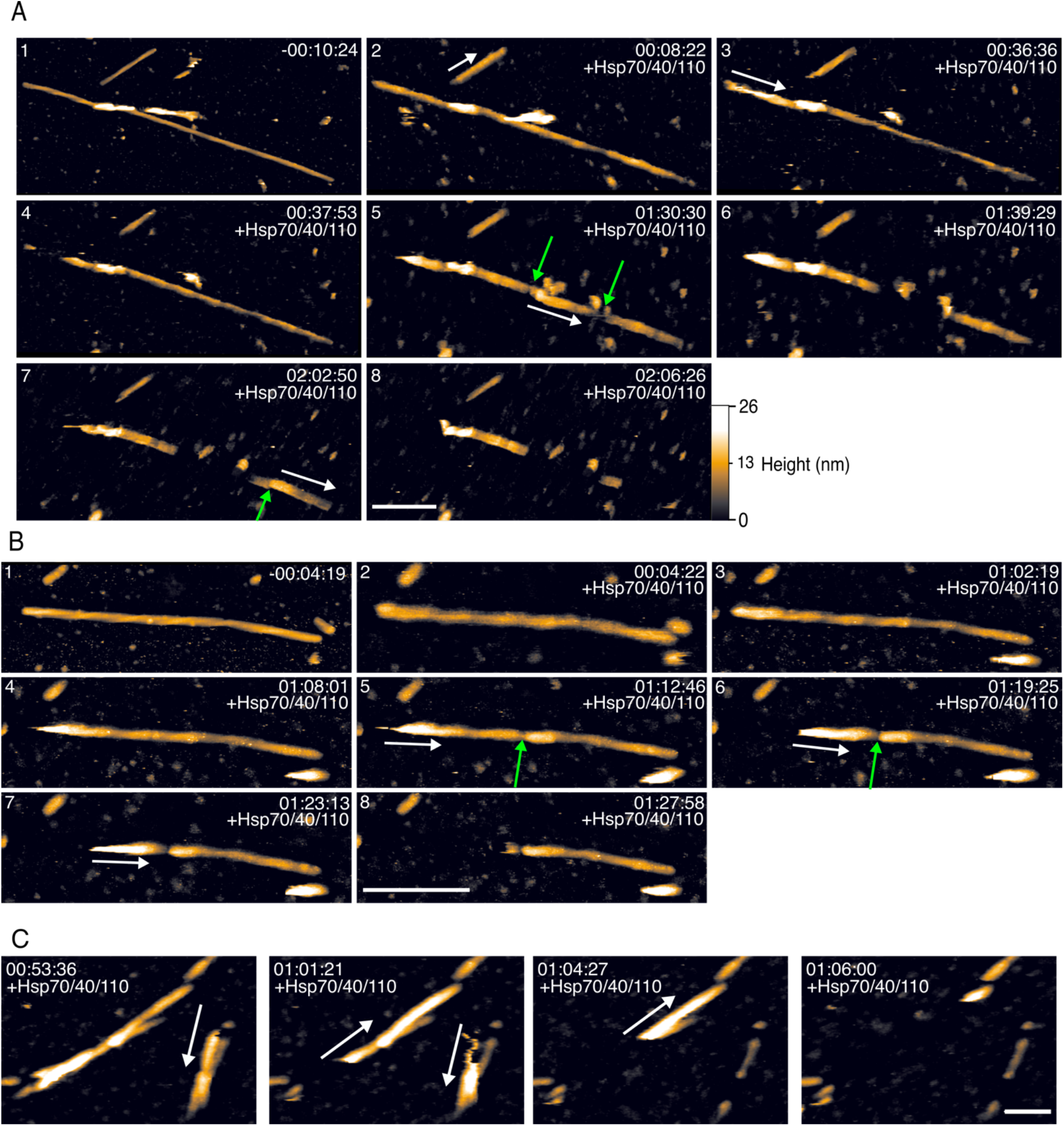
Frames from AFM videos of αSyn fibre disaggregation by Hsc70/DNAJB1/Apg2. (A) An example video series showing the disaggregation of an αSyn fibre. The time stamp is in hours:minutes:seconds, with the chaperone/ATP mixture added at time 0:00:00. Scale bar, 250 nm. The height (colour) scale is 0 (black) to 26 nm (white), see inset at bottom right, where 0 nm refers to the passivated mica substrate. (B) Video frames showing key stages of the disaggregation process. Green arrows indicate fragmentation sites. White arrows indicate the direction of depolymerisation along of fibres. Time stamps and scale bar as in (A). (C) Frames from AFM video series that show the polar disassembly of αSyn fibres. The white arrows indicate the direction of depolymerisation. Time stamps and scale bar as above. The videos are provided as supplementary videos 1 and 2.

The AFM videos showed both fragmentation and runs of progressive depolymerisation (Figure 1B). This combined fragmentation and depolymerisation is consistent with published biochemical evidence (Gao et al., 2015). Not all disaggregating fibres were fragmented. Some were disassembled by multiple runs of unidirectional depolymerisation, and others appeared unaffected by the Hsc70/DNAJB1/Apg2 machinery. Of the 22 αSyn fibres imaged in 8 videos, 5 fibres were not visibly affected by the Hsc70/DNAJB1/Apg2 disaggregase. There were no structural motifs evident by AFM that distinguished these intact fibres from those that were disassembled. Even for fibres that were disaggregated, disassembly was often incomplete (8 of 17 fibres were partially disaggregated; Figure 1A, B, Supplementary Table 1). The partial disaggregation was consistent with biochemical experiments which showed disaggregation can be incomplete (Gao et al., 2015; Wentink et al., 2020). For all observed cases of αSyn disaggregation, depolymerisation progressed in only one direction along a fibre (Figure 1C). The direction of depolymerisation was independent of AFM scanning direction, which excludes the AFM tip-sample interaction as the cause of directional depolymerisation. Instead, this directionality indicates that depolymerisation is aligned with respect to the filament structural polarity. The polar nature of αSyn fibres has been demonstrated in multiple high-resolution cryo EM reconstructions (Guerrero-Ferreira et al., 2019, 2018; B. Li et al., 2018; Y. Li et al., 2018), and has led to the hypothesis of a polar growth mechanism. Our AFM movies demonstrate that depolymerisation is polar.

In 6 out of the 8 of the videos, there was a delay of 30 – 60 minutes before any disassembly of αSyn fibres was observed (Figure 1A, B panels 2 – 3). In the other 2 videos of disaggregation, disassembly started almost immediately (5 – 10 minutes) after chaperones were injected into solution. Of note, no such delay was observed in biochemical assays of αSyn disaggregation (Gao et al., 2015; Wentink et al., 2020), implying that the disaggregation kinetics in AFM experiments may be slower due to the adhesion of the fibres to the solid support (mica). Alternatively, or additionally, fluid mixing may be less thorough in the AFM fluid cell than in solution, and may result in a slower build up of critical chaperone concentration at the fibre surface in the AFM experiment. Yet, once initiated, the chaperone-mediated disaggregation was unambiguous (Figure 1).

To ensure that observed changes in fibre structure were due to bona-fide chaperone activity, rather than due to interactions between the AFM tip and the sample, we tested the ATP-dependency of αSyn fibril disaggregation in AFM videos. Hsc70/DNAJB1/Apg2-mediated disaggregation requires ATP (Gao et al., 2015; Scior et al., 2018), whereas we expect mechanical perturbation by the AFM tip to be independent of ATP. In the absence of ATP, there was no visible disruption of αSyn fibres in the 4 AFM videos of fibres incubated with Hsc70/DNAJB1/Apg2 (Supplementary Figure 1). Therefore, the breaks and depolymerisation of αSyn fibres can be attributed to the action of the Hsc70/DNAJB1/Apg2 disaggregase, rather than to tip-sample interactions related to the AFM imaging process.

### Amyloid fibres are depolymerised in rapid processive bursts

The AFM videos of αSyn fibre disaggregation revealed that depolymerisation occurs in short, rapid bursts. In these video series, segments of fibres were typically disassembled within 2 – 3 successive video frames, where each frame lasted 1 – 2.5 minutes (Figure 2A). The lengths of these rapidly depolymerised segments varied between 85 – 375 nm with an average of 176 ± 84 nm (mean ± standard deviation, for 26 depolymerisation events from 17 fibres; initial and final fibril lengths are listed in Supplementary Table 1). We did not observe evidence of more gradual, continuous depolymerisation in any of the AFM videos.

**Figure 2.**
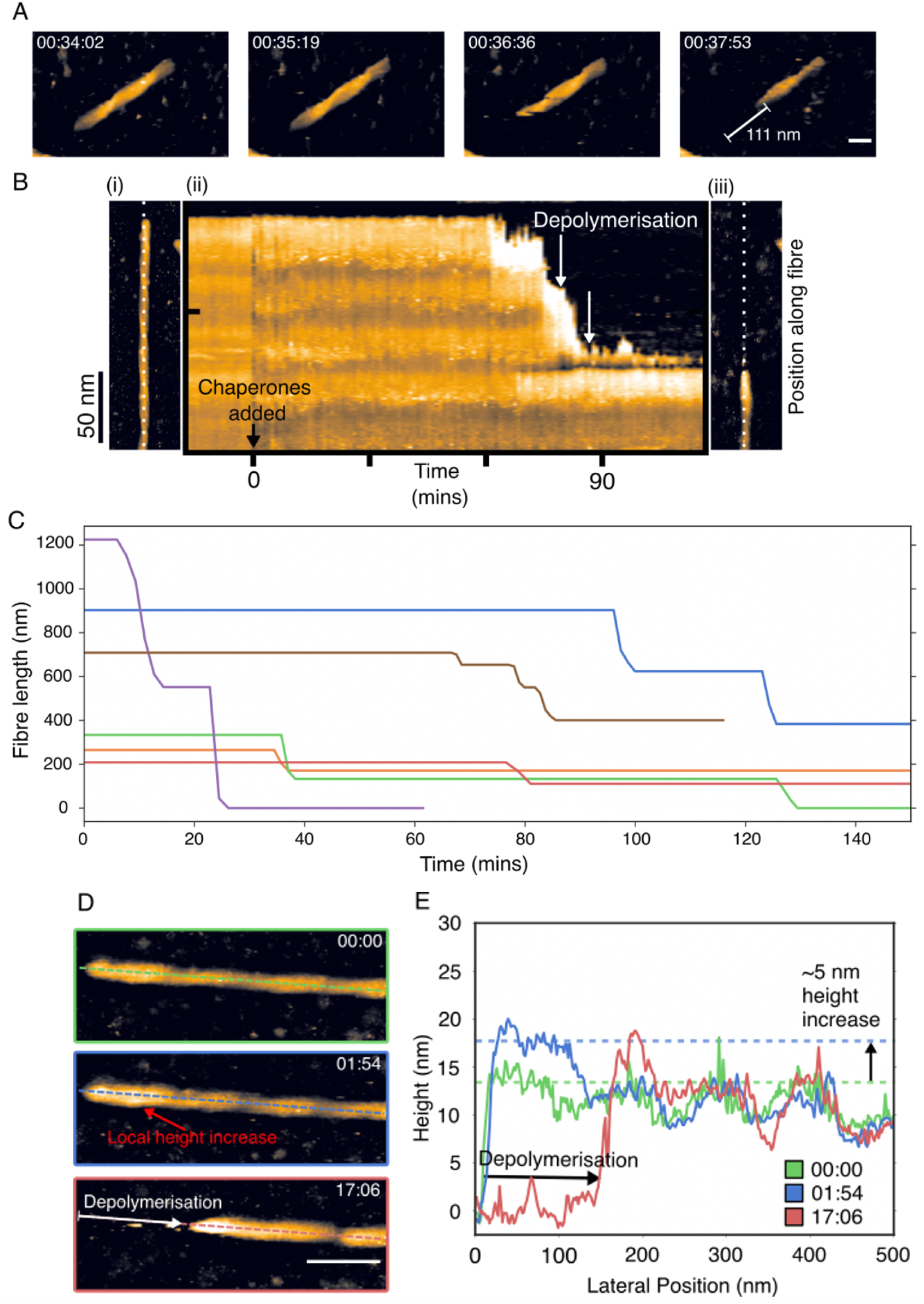
Time course of depolymerisation and morphological changes. (A) Example segment of an αSyn fibre depolymerised in a rapid “burst”, which removed 111 nm of this fibre (indicated with white marker) within 2 – 3 successive video frames (*i*.*e*., within 3 – 4 minutes). Height (colour) scale 26 nm, see Figure 4.1. (B) A kymograph showing the disaggregation of a single αSyn fibre: (i) The first frame of an AFM video series of disaggregation. (ii) Kymograph showing a height profile that is aligned to the fibre shown in A (white dotted line) as a function of time (horizontal axis of kymograph). Zero minutes is defined as the first frame recorded after the chaperones were injected into the solution. (iii) The same, now largely disaggregated, αSyn fibre as shown in A, at the end of the movie. (C) Lengths of individual fibres plotted as a function of time through disaggregation reactions as imaged by AFM. Traces are shown for a subset of 6 fibres, each in a different colour, from 4 independent experiments. (D) Disaggregation videos that show that a local height increase of αSyn fibres precedes depolymerisation. Zero time here refers to the latest frame before changes to the fibre were observed. Height (colour) scale: 21 nm, see Figure 1. Scale bar: 50 nm. (E) Height profiles along the dashed lines in D showing the changes to αSyn fibres during disaggregation.

We considered whether the burst-like nature of the depolymerisation reaction could be attributed to an artefact due to adhesion of the fibre to the mica substrate. Given that the chaperone binding and hence the depolymerisation reaction are expected to follow the helical path of the fibre structure, we would expect such a scenario to imply a correlation between the lengths of the depolymerisation bursts and the helical periodicity. The helical twist can be detected in the AFM experiments prior to chaperone injection (Figure 1A,B). Since no such correlation is observed, we rule out such an artefact (Supplementary Figure 2).

To assess the kinetics of depolymerisation, we translated the AFM videos into kymographs that displayed the height profile along a fibre as a function of time, with depolymerisation marked by a drop in measured height of the fibre thickness (8 – 12 nm) to the passivated mica substrate (∼0 nm). These kymographs showed no evidence of gradual depolymerisation and further highlighted that depolymerisation occurred in stepwise bursts (Figure 2B). We quantified this by measuring lengths of individual fibres as a function of time, through the disaggregation movies (Figure 2C). Again, this highlighted that depolymerisation occurred exclusively by discrete stepwise bursts, as shown by sharp reductions in fibre length as a function time, and that fibre lengths were stable between such bursts (Figure 2C). These rapid depolymerisation events reduced the length of a fibre at an average speed of 64 ± 36 nm/min (mean± standard deviation; median value ∼50 nm/min), whereas there were no observable changes in fibre length during the times (10 – 45 minutes) between such depolymerisation bursts (Figure 2B). The temporal resolution of the AFM videos was relatively low, typically 2 – 3 frames per depolymerisation burst. Nevertheless, it was sufficient for us to estimate the average depolymerisation speed during active bursts, corresponding to the extraction of ∼4 monomers of αSyn from a fibre per second (for a 2 protofilament fibre). This stepwise depolymerisation, with bursts of depolymerisation interspersed with static dwells, indicates that the Hsc70/DNAJB1/Apg2 machinery is not able to continuously extract αSyn monomers from one end of a fibre to the other. We therefore hypothesised that αSyn fibres need to be locally destabilised before the Hsc70/DNAJB1/Apg2 machinery can proceed with αSyn removal.

### An increase in fibre height precedes depolymerisation

Consistent with the notion of a local change preceding depolymerisation, we noticed that αSyn fibre segments displayed a characteristic morphological change preceding their disassembly.

Specifically, short (178 ± 65 nm) fibre segments underwent a uniform height increase of several nm (Figure 2D). After this increase in height, there was a lag-time of 2 – 5 minutes before the raised segment was depolymerised in a rapid burst, as described above. After depolymerisation, a similar morphological change (with up to 10 nm height increase) was observed at the newly formed blunt end of the fibre (Figure 2D, E). In contrast to these depolymerisation events, we did not observe a similar morphological change prior to fibre fragmentation. Instead, breaks in fibres appeared spontaneously with no visible changes observed to precede their formation. However, we did see a height increase in the downstream (relative to the direction of depolymerisation) segment of fibres immediately after fragmentation, which was usually followed by depolymerisation.

There are several potential explanations for the observed ∼5 nm change in fibre height. Firstly, one might attribute the height increase to a partial detachment of the fibres from the surface before depolymerisation. Such detachment would cause a gradient of increasing freedom of movement of the fibre from the detachment point, rather than a small and uniform height increase (Supplementary Figure 3). Moreover, AFM imaging of the detached region would have substantially reduced resolution due to fibre mobility. However, the experimental data show no such additional movement or loss of resolution. Alternatively, the increased height of the fibres might be caused by a change in fibre structure. However, previous negative-stain EM tomograms did not reveal structural changes that would be consistent with a uniform increase in fibre height (Gao et al., 2015). Instead, we therefore attribute the local height increase to a local recruitment of Hsc70/DNAJB1/Apg2 to fibres, forming large and extended clusters that are necessary precursors for fibre disassembly.

### Cryo-EM of αSyn fibres with bound Hsc70/DNAJB1/Apg2 shows dense clusters of chaperone binding

To investigate the molecular origin of this characteristic height increase preceding disaggregation, we visualised Hsc70/DNAJB1/ATP and Hsc70/DNAJB1/Apg2/ATP complexes on αSyn fibres by cryo-EM. Cryo-EM images of the fibres alone (Figure 3A) show ∼10 nm-thick helical filaments consistent with published data (Guerrero-Ferreira et al., 2019, 2018; Vilar et al., 2008) and with the bare fibre height we observed by AFM. In cryo-EM images of fibres in the presence of DNAJB1, Hsc70 and ATP we observe chaperone clusters scattered along the fibres (Figure 3B), and when Apg2 is added, extended regions of densely clustered chaperones are seen (Figure 3C).

**Figure 3.**
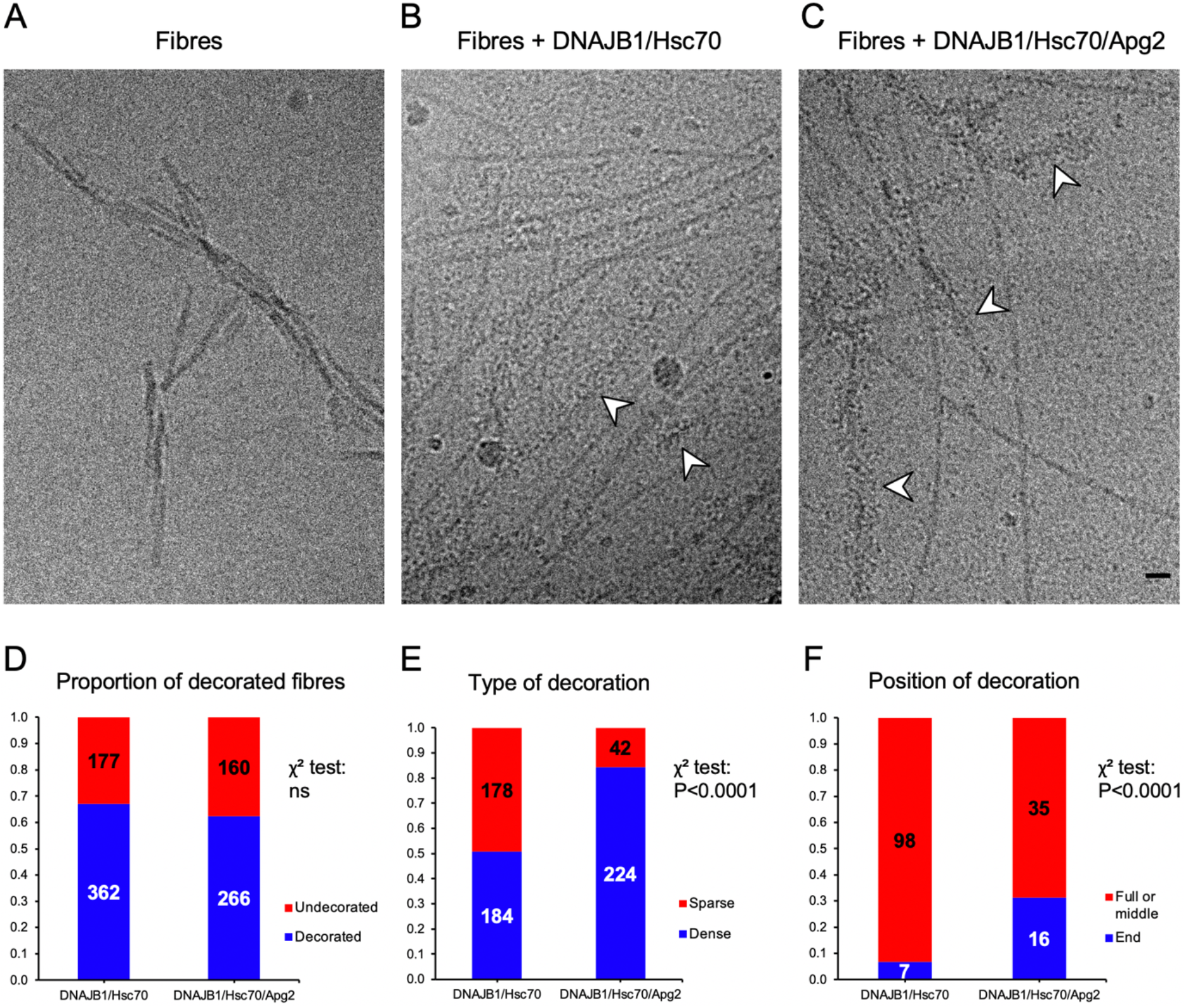
Cryo EM images and plots of fibre decoration. (A, B, C) Representative cryo EM images of αSyn fibres alone (A), αSyn fibres incubated with Hsc70/DNAJB1/ATP (B) and fibres incubated with Hsc70/DNAJB1/Apg2/ATP (C). In (B) and (C), fibres show variable binding with chaperones (white arrows) Scale bar: 30 nm. (D, E, F) Stacked bar charts of scored fibres, counting the number with chaperone decoration (D), or with dense vs sparse decoration (E). In (F), the locations of chaperone clusters along the fibres were scored. White and black numbers represent the number of counted fibres for each category. A two-sided Chi-square test was performed for each stacked bar chart. ns: Not significant.

Scoring these observations on data sets of ∼400-500 fibres counted from ∼100 micrographs (Supplementary Figure 4), we find that the relative number of fibres with bound chaperones does not significantly depend on whether Apg2 is present in the sample preparation (Figure 3D). However, the dense clusters are much more commonly seen in the presence of the full disaggregase system, suggesting that Apg2 locally enhances the recruitment of chaperones (Figure 3E). In brief, Apg2 does not affect the proportion of fibres that are decorated, but Apg2 affects the decoration itself. Consistent with the AFM observations, the denser decoration seen in the presence of Apg2 also appeared to be more often located at the fibre ends (Figure 3F).

### Hsc70 recruitment to the fibres requires both DNAJB1 and Apg2

To determine the composition of the observed decoration, we performed biochemical assays quantifying chaperone binding to the fibres. Specifically, fibres were incubated with the chaperones and ATP, followed by separation into pellet (fibre) and supernatant (unbound) fractions. Comparing the binding with and without Apg2, we found a marked increase in recruitment of Hsc70 in the presence of Apg2 (Figure 4A). Controls in the absence of fibres were also carried out to confirm that Apg2 did not trigger the aggregation of Hsc70. In addition, the DNAJB1 dependence of Hsc70 binding to fibres was verified by comparison with a mutant version lacking the J domain which is primary site of contact between Hsc70 and J-domain proteins (Figure 4B, C). Without the J domain, there was no Hsc70 recruitment detected by the binding assay in any of the conditions. In accordance with this lack of recruitment, negative stain EM of fibres incubated with ΔJ-DNAJB1, Hsc70, Apg2 and ATP do not show decoration (Supplementary Figure 5). With wild type DNAJB1, Apg2 stimulation of Hsc70 recruitment was confirmed by total internal reflection fluorescence (TIRF) microscopy. Hsc70 fluorescence intensity was higher in the presence of Apg2 (Figure 4D-F), whereas the αSyn fluorescence intensity was the same in both conditions (Figure 4F). In agreement with AFM and cryo-EM observations, TIRF intensities showed that Apg2 recruits Hsc70 preferentially to one end of each fibre (Figure 4G). This was done by comparing the Hsc70 fluorescence intensity, relative to the αSyn intensity, at the two ends of each fibre. Therefore, the binding assays show that initial recruitment of Hsc70 to the fibres requires the J domain of DNAJB1, and binding assays and TIRF confirm that Apg2 stimulates additional recruitment. The EM and TIRF microscopy show that the additional Hsc70 is recruited into dense clusters enriched at one end of the fibres.

**Figure 4.**
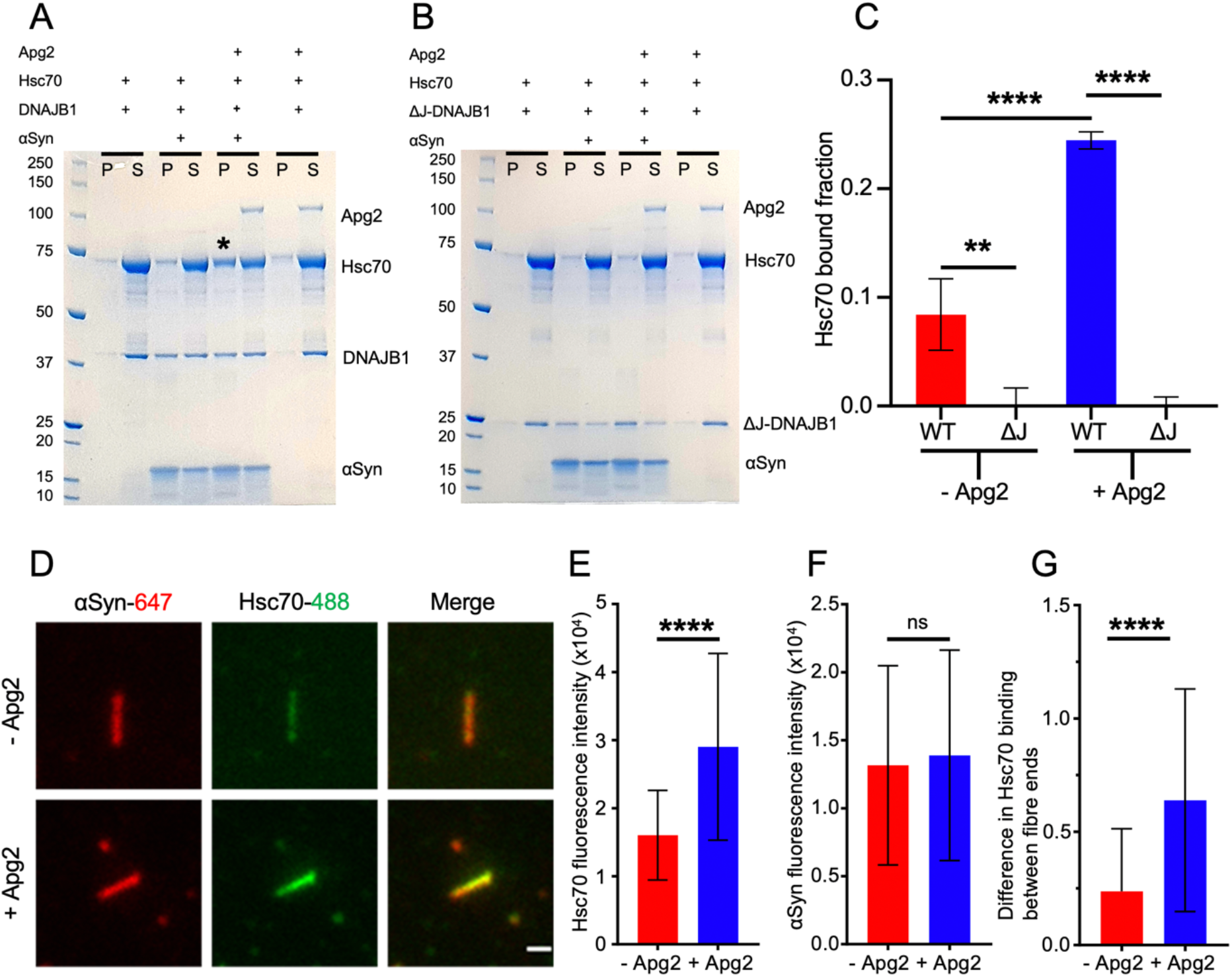
Hsc70 recruitment is greatly stimulated by Apg2. (A) Binding assay for Hsc70 binding in the presence of WT DNAJB1: control of DNAJB1/Hsc70, αSyn fibres + DNAJB1/Hsc70, αSyn fibres + DNAJB1/Hsc70/Apg2, control of DNAJB1/Hsc70/Apg2. The ‘*’ highlights the condition where an increase of Hsc70 in the bound fraction is observed. P, pellet; S, supernatant. The amount of Hsc70 recruited to the fibres is shown in the pellet fractions. (B) Binding assay for Hsc70 binding in the presence of the truncated ΔJ-DNAJB1, lacking the J domain. The gel layout is similar to the previous one with ΔJ-DNAJB1 replacing WT DNAJB1. Both Apg2 and J domain are required for enhancement of Hsc70 binding. (C) Histogram of Hsc70 bound fraction in each sample (N = 3 independent experiments, WT: WT DNAJB1, ΔJ: ΔJ-DNAJB1 mutant, -Apg2: condition with fibres + DNAJB1/Hsc70, +Apg2: condition with fibres + DNAJB1/Hsc70/Apg2). A 2-way ANOVA was performed with Tukey’s multiple comparisons test (**: P=0.003; ****: P<0.0001). (D) TIRF microscopy images of the labelled αSyn fibres and Hsc70 in the absence or in the presence of Apg2. Scale bar, 1 µm. (E, F) Quantitation of fluorescence intensity of Hsc70 (E) and αSyn (F) in the two conditions (+/- Apg2, N = 3 independent experiments, n = 65 fibres). (G) Quantitation of differences in Hsc70 binding between the two ends of the fibres, normalised to the local αSyn intensity, in the absence and presence of Apg2 (N = 3 independent experiments, n = 54 fibres). For each plot, data were tested for normality by performing a Shapiro-Wilk test. As the data did not display a normal distribution, they were analysed using a two-tailed Mann-Whitney test (****: P<0.0001, ns: not significant).

We quantified the Hsc70 recruitment observed here as a function of Apg2 concentration. By performing a binding assay with different dilutions of Apg2, we found that substoichiometric Apg2 facilitates Hsc70 recruitment, saturating at a 1:10 molar ratio of Apg2:Hsc70 (Figure 5A-C). Apg2 concentration also affected the disaggregation efficiency (Figure 5D,E). Moreover, disaggregation activity shows the same dependence on Apg2 as binding assays (Figure 5F). This shows that a key function of Apg2 in disaggregation is its concentration-dependent recruitment of Hsc70 into active assemblies.

**Figure 5.**
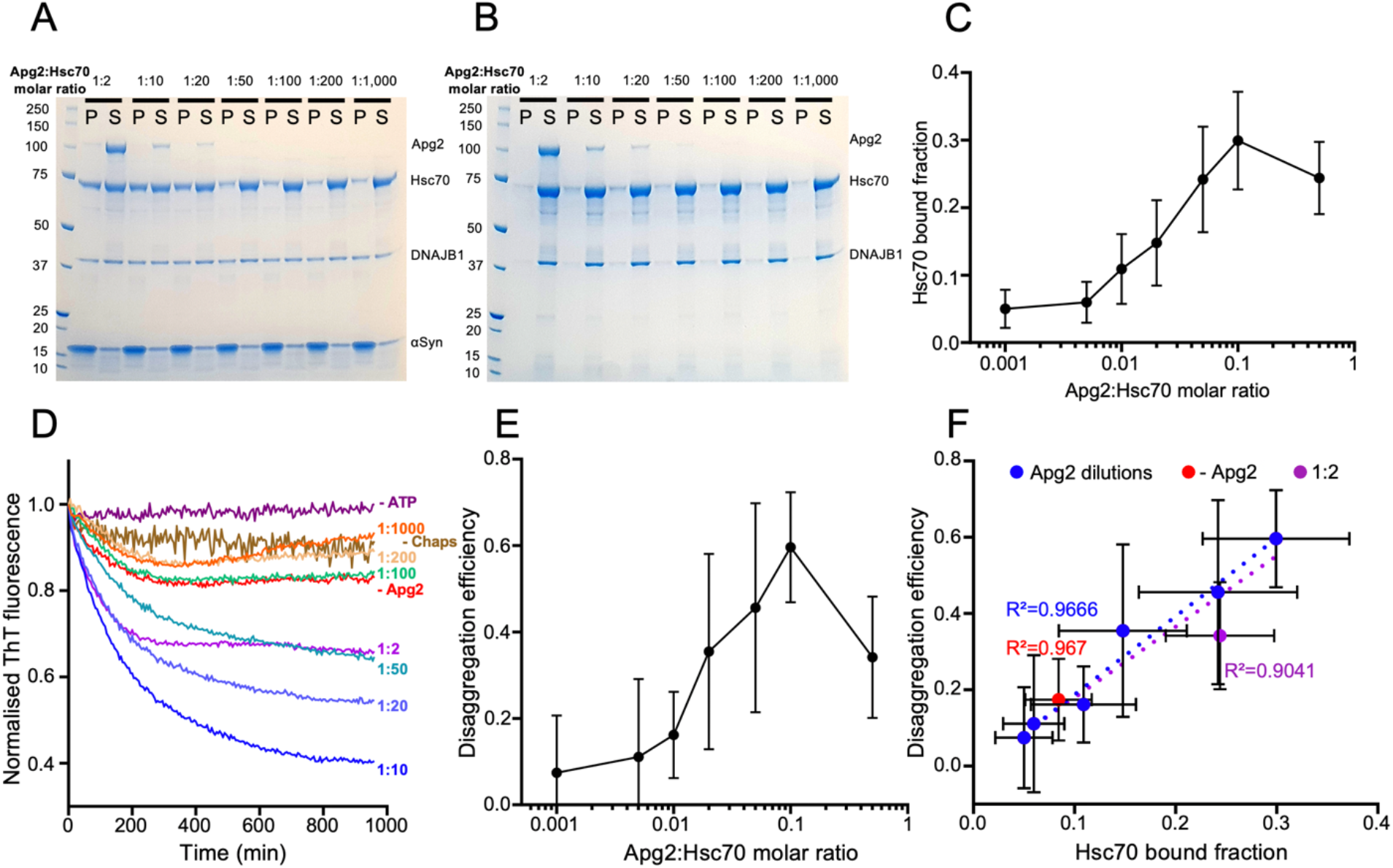
Dependence of Hsc70 recruitment and disaggregation activity on Apg2 concentration. (A) Binding assay showing the dependence of Hsc70 recruitment on Apg2. (B) Control binding assay in the absence of fibres. ATP was present in both A and B. (C) Plot showing Hsc70 binding as a function of Apg2:Hsc70 molar ratio (N = 3 independent experiments). The action of Apg2 in recruiting Hsc70 to the fibres saturates at a molar ratio of Apg2:Hsc70 of 1:10. (D) Disaggregation activity measured by Thioflavin T fluorescence shows the same dependence on Apg2 as binding assays. High Apg2 (1:2 molar ratio) makes disaggregation less efficient, possibly by prematurely releasing Hsc70. (E) Quantification of the disaggregation efficiency as a function of Apg2:Hsc70 molar ratio (N = 3 independent experiments). The disaggregation efficiency was determined by taking the values from the final hour of the assay curves in (D). (F), Plot showing the linear correlation between disaggregation activity and Hsc70 binding. Including the data point without Apg2 did not change the correlation coefficient, which remained at 0.967). In contrast, including the highest Apg2 concentration (Apg2:Hsc70 molar ratio of 1:2) reduced the correlation coefficient to 0.904.

### Cryo electron tomography shows flexible but densely packed clusters of chaperones along α-synuclein fibres

Electron tomography of the decorated fibres shows clusters of elongated complexes, flexibly extending from the fibre surface, either in patches or more continuous arrays (Figure 6). In the absence of Apg2, the binding is less dense and there are fewer and shorter clusters, with more unbound chaperones in the background (Figure 6A). With Apg2, there are extended regions with densely clustered chaperones (Figure 6B). These regions of continuous binding follow a roughly helical track, as expected from the underlying helical twist of the fibres (supplementary video of tomogram).

**Figure 6.**
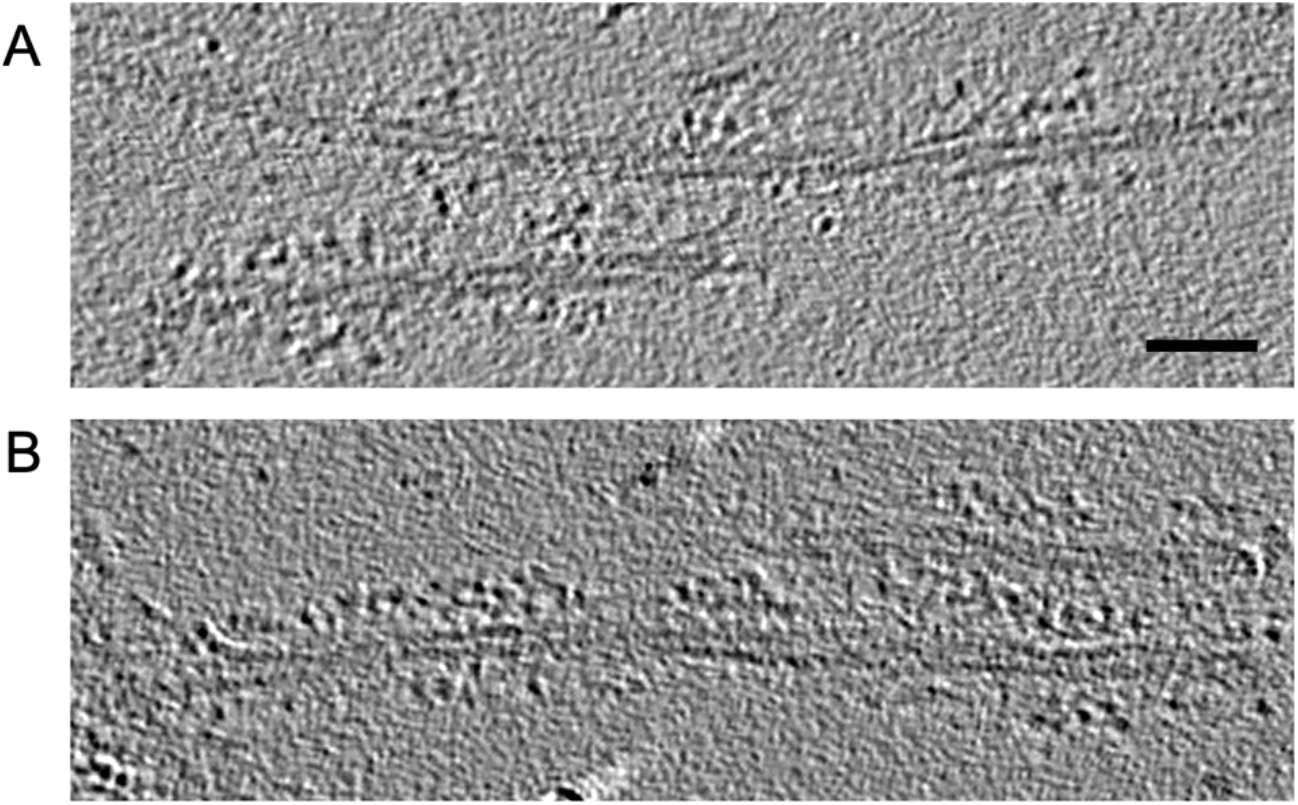
Cryo-electron tomography of fibre-chaperone complexes. (A) Tomogram section of fibres incubated with Hsc70, DNAJB1 and ATP. Small clusters are interspersed with sparse decoration. (B) Fibres incubated with Hsc70, DNAJB1, Apg2 and ATP, showing extended stretches of densely clustered chaperones. The two-protofilament structure of the fibres is evident, but the flexible αSyn N- and C-terminal tails that contain the chaperone binding sites are not discernible. Scale bar, 30 nm.

The cryo-EM and tomography show that the chaperones extend away from the fibres, qualitatively consistent with the height increase in the AFM images. For such long and flexible complexes, we expect the AFM measurement to show reduced dimensions because molecular compression and/or motion due to the force exerted by the AFM tip, which explains the difference between the 10-20 nm extension in the EM data and the ∼5 nm height measured in the AFM experiments. In brief, the cryo-EM and AFM data are consistent in showing local, dense recruitment of chaperones, which is required for localised bursts of depolymerisation.

The EM, TIRF, binding and disaggregation assays establish that Hsc70 recruitment to the fibres is strongly stimulated by Apg2. This suggests that Apg2 is responsible for the increased chaperone clustering, even though it is a relatively minor component of the complex (At 1/10^th^ the concentration of Hsc70, it is not visible in the Coomassie stained pellet fractions, Figure 4). The effect of Apg2 saturated at a 1:10 molar ratio of Apg2:Hsc70 (Figure 5). There was no obvious change in DNAJB1 binding after Apg2 addition, despite the clear increase in Hsc70 binding. This does not preclude an essential role for DNAJB1: no Hsc70 binding was observed when the DNAJB1 was substituted by truncated DNAJB1 lacking the J-domain and G/F linker (Figure 4B, C and negative stain images in Supplementary Figure 5). Our findings show that the large chaperone clusters seen on Hsc70/DNAJB1/Apg2 incubated fibres are composed predominantly of densely packed Hsc70 and DNAJB1. The binding of DNAJB1 was previously estimated as one dimer every 5 nm (Gao et al, 2015), and the spacing of the globular features in Figure 6B is roughly 6 nm. This suggests that Hsc70 and DNAJB1 are similar in mass/unit length of fibre, within a factor of ∼2.

## Discussion

In this study, we have used AFM to directly visualise disassembly of αSyn fibres by the Hsc70/Apg2/DNAJB1 disaggregase in real time. In agreement with previous work (Gao et al., 2015), the AFM videos show that Hsc70-mediated disaggregation of αSyn fibres proceeds by combined fragmentation and depolymerisation. Additionally, the videos provide further insight into the disaggregation mechanism: αSyn fibres were disassembled by polar depolymerisation that occurred in rapid bursts, in which short fibre segments were disassembled within 2 – 5 minutes.

These depolymerisation bursts are facilitated by stretches of densely clustered chaperones, as observed by cryo EM, and appearing as a local height increase along fibre segments in the AFM images just prior to disaggregation bursts. The clusters consist largely of Hsc70 and DNAJB1, but substoichiometric amounts of Apg2 are required to stimulate further Hsc70 recruitment and to create the extended, dense clusters.

Previous studies have shown that Hsc70 clustering is necessary for amyloid disaggregation, but the FRET analysis in those studies did not reveal Apg2-stimulated additional recruitment of Hsc70 to the fibrils (Faust et al., 2020; Wentink et al., 2020). We have directly visualised these clusters by cryo electron tomography, showing that Hsc70/DNAJB1/Apg2 forms densely packed chaperone clusters that stretch over extended (>∼100 nm) lengths along the αSyn fibres.

Earlier negative stain EM tomography showed that DNAJB1 binds all along the fibres, and that, individually, Hsc70 and Apg2 bind irregularly to the fibres (Gao et al, 2015). It has been shown that the JDP plays a crucial role in recruiting Hsc70 to the fibres, with a specific requirement for release of the auto-inhibitory interaction in DNAJB1 (Faust et al, 2020). In addition, Hsc70 clustering driven by specific Apg2 NEF activity is important for disaggregation (Wentink et al, 2020). Our analysis of the complexes reveals an important, dose-dependent effect of Apg2 in recruiting additional Hsc70 into dense clusters on the fibres, and in stimulating the bursts of disaggregation. It is surprising that a recycling factor (NEF), which is expected to release Hsc70 from its substrate, increases total Hsc70 bound to the fibrils.

Our AFM videos suggest that chaperone clustering is a necessary and direct precursor to processive depolymerisation of the fibres, consistent with the previous demonstration that cluster formation is required for fibre disassembly. Interestingly, we observe such clusters predominantly at fibre ends (or covering an entire short fibre), in both cryo EM and AFM images. In combination with the observed unidirectional depolymerisation, this suggests that the Hsc70/DNAJB1/Apg2 machinery preferentially pre-assembles at one end of the polar αSyn fibres (Guerrero-Ferreira et al., 2019, 2018; B. Li et al., 2018) for rapid, processive depolymerisation.

Based on these observations, we propose a model for αSyn fibre depolymerisation, initiated when substoichiometric amounts of Apg2 stimulate binding of Hsc70 into extended clusters, preferentially at fibre ends (Figure 7). We speculate that these large clusters are required to provide sufficient pulling force to locally destabilise αSyn fibres, and that this destabilisation primes a fibre segment for subsequent rapid and cooperative extraction of αSyn monomers by the Hsc70/DNAJB1/Apg2 complex. Once initiated, this depolymerisation then proceeds rapidly along the whole decorated segment. This process appears to be the reverse of the assembly mechanism of amyloid fibres: previous studies have reported polar assembly (fibre growth faster in one direction along the fibre than the other) of Aβ_1-42_ fibres (Watanabe-Nakayama et al., 2016; Young et al., 2017), the functional bacterial amyloid curli (Sleutel et al., 2017) and for subpopulations of the Sup35 yeast prion (DePace and Weissman, 2002). The chaperone-mediated polar depolymerisation reported here could represent a reversal of this assembly mechanism, in which depolymerisation is energetically favoured in one direction, and chaperone clustering is nucleated at the preferred end of the fibres. Furthermore, one hypothesis for amyloid nucleation is that the process is initiated by oligomer formation followed by a conformational rearrangement that results in the amyloid structure (Chatani and Yamamoto, 2018; Nors Pedersen et al., 2015). Again, we speculate that the disassembly of amyloid could represent the reversal of this process, whereby a massive local recruitment of the Hsp70 disaggregase machinery is required to provide sufficient force to overcome the activation energy for partial amyloid unfolding, forming a more disordered aggregate which can then be rapidly disassembled.

**Figure 7.**
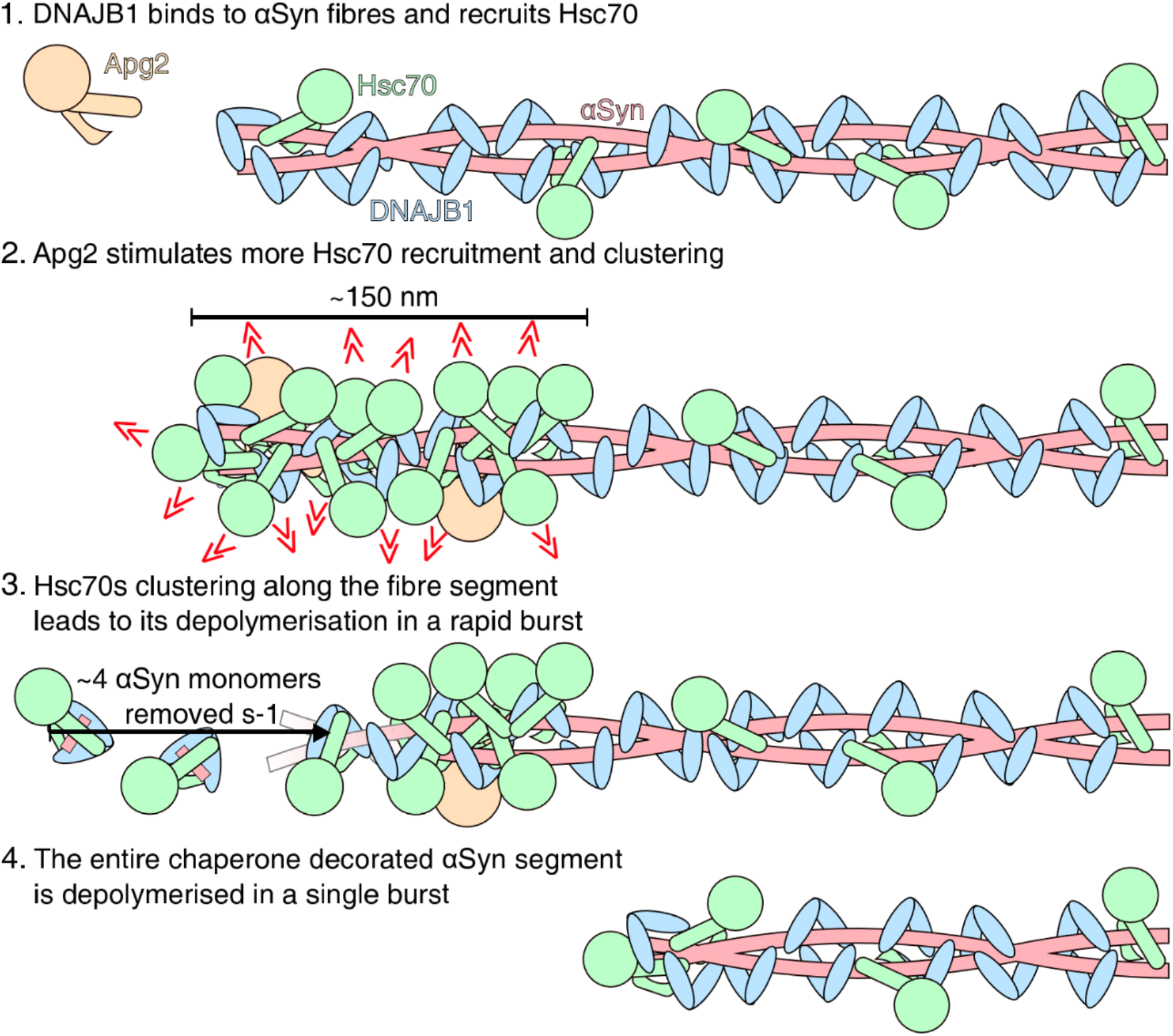
Proposed model of αSyn fibre depolymerisation. Apg2 catalyses the assembly of densely packed clusters of Hsc70, priming the rapid bursts of disaggregation.

In summary, we have presented the first views of the dense arrays of chaperones that bind along segments of αSyn fibres to form tracks that subsequently trigger bursts of rapid, unidirectional disassembly. It remains a future challenge to understand the dynamics of the Hsc70 system during these bursts.

## Materials and Methods

### Hsc70 and Apg2 expression and purification

Hsc70 and Apg2 were expressed in *E. coli* Rosetta™ DE3 cells with an N-terminal His_6_-Smt3 tag and purified by nickel affinity chromatography. Protein expression was induced using IPTG. Cell lines were grown in LB media, at 37 °C, to an OD_600_ density of 0.6 – 0.8 and 0.5 mM IPTG was added to the cell cultures. After induction, cell cultures were incubated for a further 3 hours before cells were harvested by centrifugation (4,200 g, 20 minutes).

Hsc70 and Apg2 were purified by identical protocols. Pellets from respective cell lines were resuspended in Hsc70/Apg2 lysis buffer (Supplementary Table S2), with an EDTA free protease inhibitor tablet (Roche), and lysed by cell disruption. Cell debris and insoluble protein was removed by centrifugation (48,000g, 60 minutes) before the soluble fraction was collected and incubated with Ni-IDA beads for 2 hours at 4 °C. Contaminant proteins were removed from the column in 3 successive washes: first using Hsp70/110 wash 1 buffer, followed by Hsp70/110 wash 2 buffer, and finally 2 washes with ATP wash buffer (Supplementary Table S2). Hsc70 and Apg2 were isolated using on column cleavage in which the resin was incubated with Ulp1 protease at 4 °C overnight.

After this, the flow through was collected and the column was washed twice with Hsp70/110 lysis buffer and these fractions were collected. Hsc70 samples were concentrated to 85 µM and Apg2 samples were concentrated to 25 µM. Samples were aliquoted, flash frozen in liquid nitrogen and stored at -80°C. The T111C Hsc70 (T111C, C267A, C574A, C603A) mutant was expressed and purified as described above. DTT was replaced by TCEP in all the buffers.

### DNAJB1 expression and purification

DNAJB1 was expressed using the same construct design as for Hsc70 and Apg2 (with an N-terminal His6-Smt3 tag) and with the same protocols. DNAJB1 was purified using nickel affinity purification and size exclusion chromatography (SEC). Cell lines for DNAJB1 were lysed by cell disruption in DNAJ lysis buffer, the resulting sample was centrifuged (48,000 g, 60 minutes) and the soluble fraction was collected and used for purification.

For DNAJB1 purification, the construct was bound to an Ni-IDA column as described above for Hsp70/110 and contaminant proteins were removed from the column by two successive wash steps, using DNAJ wash-1 buffer followed by DNAJ wash-2 buffer (Supplementary Table S2). DNAJB1-His6-Smt3 was eluted with DNAJ elution buffer (Supplementary Table S2) and collected. The His6-Smt3 tag was removed using Ulp1 during overnight dialysis into DNAJ lysis buffer (Supplementary Table S2). After the tag was cleaved, the solution containing DNAJB1 was reapplied to a Ni-IDA column to remove Ulp1, the His6-Smt3 tag and uncleaved DNAJB1-His6-Smt3. The flow through and two column washes with DNAJ lysis buffer fractions were collected and further purified by SEC.

SEC was performed using a Superdex 200 Increase 10/300 gel filtration column (GE Healthcare) using DNAJ SEC buffer (Supplementary Table S2). Pure fractions were collected and concentrated to ∼250 µM. Samples aliquoted, flash frozen in liquid nitrogen and stored at -80 °C.

The purified truncated DNAJB1 mutant lacking the J-domain and the G/F linker (ΔJ-DNAJB1) was kindly provided by Rina Rosenzweig (Weizmann Institute). Briefly, the mutant was purified by ion metal affinity chromatography (IMAC, Nickel), dialysed and cleaved with TEV protease, subjected to reverse IMAC and then subjected to SEC. The protein was stored in 25 mM HEPES pH 7.5, 200 mM KCl, 10 mM MgCl_2_, 2 mM DTT.

### αSyn purification

αSyn was expressed as a native untagged sequence in DE3 Star E. coli cell lines (Thermo Fisher Scientific, USA) by identical protocols to those described above for Hsp70/110. Cell lines were lysed by cell disruption in αSyn lysis buffer (Supplementary Table S2), the resulting sample was centrifuged (48,000g, 60 minutes) and the soluble fraction was collected. This soluble fraction was boiled for 20 minutes to aggregate *E. coli* cellular proteins and centrifuged (21,000g, 30 minutes) to pellet the aggregates. *E. coli* nucleic acids were precipitated by adding 30 mg/mL streptomycin sulphate to the soluble fraction which was then incubated for 30 minutes at 4°C. The resulting solution was centrifuged (20 min, 20,000g) to pellet the precipitated nucleic acids and the supernatant was collected. αSyn was precipitated by adding 400 mg/mL ammonium sulphate and incubating the resulting mix at 4°C for 30 minutes. The resulting sample was centrifuged (20 min, 20,000g), the supernatant was discarded, and the pellet, containing αSyn, was resuspended in TBS buffer (Supplementary Table S2) and dialysed overnight into deionised water.

The dialysed sample was purified by ion exchange chromatography with a HiTrap Q HP column using αSyn buffer A and B followed by further purification by SEC using αSyn SEC buffer (Supplementary Table S2). Pure fractions were collected and used to form amyloid fibres.

The S9C αSyn mutant was expressed and purified as described above. TCEP was added to a final concentration of 2 mM in the purification buffers.

### αSyn fibrillation reaction

We produced αSyn fibres as previously described (Gao et al., 2015): 1 mL of 200 µM monomeric αSyn was incubated in fibrillation buffer (Supplementary Table S2) in a 1.5 mL Protein LoBind Eppendorf tube (Eppendorf, Germany) on an orbital shaker (PCMT Thermoshaker Grant-bio) at 1000 rpm, 37°C for 1 week. We confirmed amyloid fibres had formed, rather than amorphous or small aggregates, by negative stain electron microscopy. S9C:WT (1:2, molar ratio) αSyn fibres were prepared as described above. TCEP was added to a final concentration of 2 mM in the fibrillation buffer.

### AFM Sample Preparation

αSyn fibres were immobilised on freshly cleaved mica disks (diameter 12 mm) by non-specific adsorption: 20 – 50 μM αSyn fibres (monomer concentration) were added to a 20 µL droplet of HEPES buffer (Supplementary Table S2) on a mica surface. This sample was incubated at room temperature for 30 minutes after which the surface was washed 3 times with 50 µL HEPES buffer.

We used a poly(ethylene glycol)-block-poly(L)-lysine (PEG-PLL) (Alamanda Polymers, USA) copolymer composed of 22 repeats of polyethylene-glycol (PEG) and 10 repeats of poly-L-lysine (PLL) to reduce non-specific binding of chaperones to the mica surface (Akpinar et al., 2019). A layer of PEG-PLL was added to the remaining mica surface after αSyn adsorption by adding 0.5 µL 1 mg/mL PEG-PLL to the 20 µL HEPES buffer droplet. This was incubated for 1 minute before the surface was thoroughly washed 10 times with 60 µL HKMD buffer (Supplementary Table S2).

In preparation for disaggregation reactions, the buffer was exchanged by 3 washes with 50 μL disaggregation buffer (table S2 Supplementary Table S2) or with HKMD buffer for controls without ATP. After buffer exchange, 40 µL disaggregation buffer (HKMD for -ATP controls) was added to the sample droplet. A 20 µL droplet of disaggregation buffer (HKMD buffer for -ATP controls) was added to the cantilever loaded on to the Z-scanner (FastScan) or fluid cell (Multimode 8). The final sample volume was ∼80 µL.

### AFM Imaging of Disaggregation

All AFM imaging was performed in liquid using Peakforce Tapping with either a Dimension FastScan microscope (images presented in Figures 1A, B, Ci and Figures 2 and 3) or a Multimode 8 (images presented in Figure 1Cii). FastScan D cantilevers (Bruker) were used with the Dimension FastScan microscope and PeakForce HIRS-F-B cantilevers (Bruker) were used with the Multimode 8 microscope. FastScan D cantilevers were operated using a drive frequency of 8 kHz and PeakForce HIRS-F-B cantilevers were operated at 4 kHz drive frequency. The areas imaged were 0.8 × 0.25 – 4 × 2 µm^2^ in size and recorded at 3.5 Hz (FastScan) or 1.75 Hz (Multimode 8) line rate. Images were 512 × 256, 512 × 172 or 384 × 172 pixels in size.

During AFM movie acquisition, disaggregation was initiated by retracting the cantilever tip ∼100 nm from the surface and injecting 0.75 – 1 µM Hsc70, 0.37 – 0.5 µM DNAJB1 and 0.07 – 0.1 µM Apg2 (always at a Hsc70:DNAJB1:Apg2 molar ratio of 1:0.5:0.1) in disaggregation buffer (HKMD buffer for -ATP controls) into the sample droplet. AFM imaging was continued within 1 minute of injection of chaperones. The same area of αSyn fibres was imaged for >2 hours at either at room temperature or at 30 °C.

### AFM Image Processing

AFM images were processed using the Gwyddion 2.52 software package (Nečas and Klapetek, 2012). Raw data was initially corrected using plane-subtraction to remove sample tilt. After this, images were corrected for systematic height errors between fast-scan (*x*-axis) line profiles using median background subtraction for each line. Additional distortions were removed by 2^nd^ order polynomial subtraction, using a polynomial calculated using only pixels corresponding to the mica/PEG-PLL surface. Higher-frequency noise was reduced by a gaussian filter with a full width at half maximum of 2 – 3 pixels, corresponding to 3 – 8 nm. The height values in the image were offset such that the mean pixel value was set to zero. The colour scale was set to an appropriate range (listed in the Figure legend for each image) and images were saved in TIFF format.

Images corresponding to a video series were opened as an image sequence using the ImageJ Fiji software package (Schindelin et al., 2012). Images in each time-series were aligned as rigid bodies using the Fiji module “StackReg” (Thevenaz et al., 1998) to correct for lateral drift between frames. Aligned movies were then cropped and saved as individual TIFF images and as AVI movies for analysis.

### Cryo EM sample preparation

αSyn amyloid fibres were sonicated for 30 minutes using a Branson 1800 ultrasonic cleaner to produce dispersed fragments of αSyn fibres. The suspension was diluted to 6 µM αSyn monomer concentration and mixed, when indicated, with 6 µM Hsc70, 3 µM DNAJB1 and 0.6 µM Apg2, and incubated for 1 hour at 30°C in disaggregation buffer (or HKMD buffer for the control without chaperones). 4 µL of this preparation were applied to negatively glow discharged holey carbon C-flat grids (CF-1.2/1.3-4C) (Protochips, USA), back blotted and plunge frozen in liquid ethane using a Leica EM GP2 (Leica Microsystems, Germany).

For the cryo-electron tomography experiments, similar samples were prepared. The sample was applied to glow discharged holey carbon grids which was then back blotted before adding 3 µL of 10 nm Protein-A-coated gold (EMS, USA) as fiducial markers for 3D reconstruction of tilt series. The grids were then back blotted a second time before plunge freezing.

### Negative Stain sample preparation

αSyn fibres (10 µM monomer) were mixed, when indicated, with Hsc70 (10 µM), Apg2 (1 µM), and WT or ΔJ-DNAJB1(5 µM monomer) for 1 hour at 30°C in the disaggregation buffer (or HKMD buffer for the control without chaperones). These samples were applied to glow discharged continuous carbon film-coated, 300 mesh copper grids (EMS) and blotted after 1 minute. The grid was then stained with 2% (w/v) uranyl formate and immediately blotted 3 times.

### EM Data Collection

Images displayed in Figure 4 and negative stain images in supplementary Figure 4 were acquired using a Tecnai TF20 electron microscope (Thermo Fisher, USA) equipped with FEG source operating at 200 keV. Images were recorded with a DE-20 direct electron detector (Direct Electron, USA) at a magnification corresponding to 1.83 Å/pixel using a total dose of ∼30 e^-^/Å^2^.

Images of the sample without Apg2 (Supplementary Figure 4) were acquired at the Electron Bio-Imaging Centre (eBIC) at Diamond on a post-GIF K3 detector (Gatan, USA) operating in super resolution counting mode using a Titan Krios microscope (Thermo Fisher, USA) operating at 300 keV. A Bioquantum energy filter (Gatan, USA) was used with a slit width of 20 eV. 50-frame movies were collected using an exposure time of 88 ms and a dose of 1.084 e^-^/Å^2^ per frame resulting in a total dose of 54.22 e^-^/Å^2^ per movie with a super resolution pixel size of 0.53 Å. 2,424 movies were collected using a 1.5 - 3.3 μm defocus range.

Images of the sample with Apg2 used for statistical analysis (Supplementary Figure 4) were acquired at Birkbeck College, University of London on a post-GIF K3 detector (Gatan, USA) operating in super resolution counting mode using a Titan Krios microscope (Thermo Fisher, USA) operating at 300 keV. A Bioquantum energy filter (Gatan, USA) was used with a slit width of 20 eV. 50-frame movies were collected using an exposure time of 50 ms and a dose of 1.0218 e^-^/Å^2^ per frame resulting in a total dose of 51.09 e^-^/Å^2^ per movie with a super resolution pixel size of 0.5335 Å. 4,039 movies were collected using a 1.5 - 3.3 μm defocus range.

Movies were gain corrected, binned by a factor of 2, motion corrected, and dose weighted using the CPU implementation of MotionCor2 in RELION (Zheng et al., 2017; Zivanov et al., 2018). Images were then denoised using the U-net (small) model in TOPAZ-Denoise (Bepler et al., 2020).

Tilt series of both conditions (without or with Apg2) were acquired on the Titan Krios at Birkbeck College using the same detector, energy filter and operating voltage. 10-frame movies were acquired per tilt angle. Tilt series were acquired with SerialEM (Mastronarde, 2005) from -60° to +60° with a 3° increment using a dose symmetric tilt-scheme with a super resolution pixel size of 1.065 Å and a 2 - 5 μm defocus range. The total dose was around 150 e^-^/ Å^2^. 30 and 23 tilt series were collected of the conditions without or with Apg2, respectively. Frames underwent whole-frame alignment in MotionCor2 version (Zheng et al., 2017) and were dose weighted before reconstructing the tomograms using IMOD version 4.9.0 (Kremer et al., 1996). Tomograms were binned by 4 in X and Y. The tomogram of the sample with Apg2 was CTF-corrected in IMOD.

### Sedimentation Assay

All chaperone aliquots (Hsc70, DNAJB1, Apg2) were centrifuged at 17,000 g for 30 minutes at 4°C. αSyn amyloid fibres were sonicated for 15 minutes at high frequency using a CPX 2800 Bransonic Ultrasonic bath (Branson). αSyn amyloid fibres (40 µM, monomer concentration) were incubated with Hsc70 (8 µM), DNAJB1 or ΔJ-DNAJB1 (4 µM), and Apg2 (different concentrations as indicated) in the disaggregation buffer for 1 hour at 30°C. The samples were then centrifuged for 45 minutes at 17,000 g to separate the pellet, containing the fibres and bound chaperones, from the supernatant with unbound chaperones. The supernatant was then collected and both fractions were incubated in 4X NuPAGE LDS Sample buffer (Thermo Fisher Scientific) for at least 30 minutes at 90°C. Samples were then loaded on BOLT 4-12% Bis-Tris gels (Thermo Fisher Scientific) and proteins were separated by SDS-PAGE. At least 3 independent experiments were performed for each condition. The statistical analyses were performed in Prism 8 (GraphPad). Data were analysed using a 2-way ANOVA test with Tukey’s multiple comparisons test.

### Thioflavin T Assay

In order to keep the αSyn amyloid fibres well dispersed and accessible to the chaperones, they were sonicated for 15 minutes at high frequency using a CPX 2800 Bransonic Ultrasonic bath (Branson). Thioflavin T (ThT) fluorescence was recorded every 5 minutes for 16 hours on a FLUOstar Omega plate-reader (BMG LABTECH, excitation: 440 nm, emission: 482 nm) to monitor the disaggregation of αSyn amyloid fibres. αSyn amyloid fibres (2 µM, monomer concentration) were incubated with Hsc70 (4µM), DNAJB1 (2 µM), and Apg2 (different concentrations as indicated) in the disaggregation buffer and 15 µM ThT. Background ThT fluorescence of chaperones and buffer was subtracted and ThT fluorescence was normalised to the fluorescence intensity of the first time point (t = 0 min). At least 3 independent experiments were performed for each condition. The disaggregation efficiency was calculated by averaging the values of the final hour for each experiment. The plots of normalised intensity over time as well as the linear regression showing the disaggregation efficiency as a function of Hsc70 bound fraction were plotted and calculated in Microsoft Excel (for Windows 365, version 16.0.13127.21624).

### TIRF Microscopy Sample Preparation

S9C:WT (1:2) αSyn amyloid fibres were incubated with Alexa Fluor 647 maleimide dye (10-fold molar excess, Thermo Fisher Scientific) and Biotin-X, SE dye (2.5-fold molar excess, Thermo Fisher Scientific) for at least 2 hours at room temperature in the dark in HKMT buffer. T111C Hsc70 was incubated with Alexa Fluor 488 maleimide dye (10-fold molar excess, Thermo Fisher Scientific) for at least 2 hours at room temperature in the dark in HKMT buffer. Labelled Hsc70 was then buffer exchanged using PD SpinTrap G-25 column (Cytiva) pre-equilibrated in HKMD buffer. Labelled amyloid fibres (1 µM) were incubated with labelled Hsc70 (2 µM), DNAJB1 (1 µM) and Apg2 (0.2 µM, if indicated) in disaggregation buffer for 1 hour at 30°C in the dark.

### TIRF Microscopy Imaging and Analysis

Chambers were prepared using glass slides, biotin-PEG coverslips, and double-sided tape. Chambers were passivated with 0.5% BSA for at least 10 minutes, washed twice in HKMD buffer, incubated twice in 0.5 mg/mL neutravidin for 2 min, washed three times in HKMD buffer, incubated with the sample for 10 minutes, and washed three times in HKMD buffer. Samples were imaged on an Eclipse Ti-E inverted microscope with a CFI Apo TIRF 1.49 N.A. oil objective, Perfect Focus System, H-TIRF module, LU-N4 laser unit (Nikon) and a quad band filter set (Chroma), described in Toropova et al., 2017. Frames were recorded on an iXon DU888 Ultra EMCCD camera (Andor), controlled with NIS-Elements AR Software (Nikon) with an exposure time of 100 ms.

To determine the overall fluorescence of the fibres, Hsc70 and αSyn fluorescence intensities were measured in ImageJ FIJI software package by determining the plot profile of 65 fibres for each condition. The minimum value for each plot profile was considered as background and subtracted. At least 3 independent experiments were performed for each condition. The statistical analysis were performed in Prism 8 (GraphPad). Data were first analysed by running a Shapiro-Wilk test to check their normality. A two-tailed Mann Whitney test was then performed as the data did not have a normal distribution.

To evaluate the distribution of bound Hsc70, Hsc70 and αSyn fluorescence intensities were measured along the fibres. The intensities in the first and last 3 pixels of each fibre were compared: the ratio of Hsc70 intensity divided by αSyn intensity was calculated at each end and the lower ratio was then subtracted from the higher one. The division by αSyn intensities served to minimise the influence of variations in fibre distance from the glass surface. The difference between the two ratios was plotted in Figure 4G. 54 fibres from 3 independent experiments were measured.

Statistical analyses were performed as described above using Prism 8 (Shapiro-Wilk test followed by a two-tailed Mann Whitney test as the data did not have a normal distribution).

## Acknowledgements

All the cryo-EM samples and most of the cryo-EM data (all but the 2,424 movies of the sample without Apg2) for this investigation were collected at the ISMB EM facility at Birkbeck College, University of London with financial support from the Wellcome Trust (202679/Z/16/Z and 206166/Z/17/Z). We acknowledge Diamond Light Source for access and support of the cryo-EM facilities at the UK’s national Electron Bio-imaging Centre (eBIC) [under proposal EM 20287], funded by the Wellcome Trust, MRC and BBRSC. We thank Natasha Lukoyanova for Krios data collection, and for comments on the manuscript. The TIRF microscope was supported by grants from the Wellcome Trust (217186/Z/19/Z and 210585/Z/18/Z) and the Royal Society (RG170260). We thank the Baden-Württemberg Stiftung (BWST-INTSFIII-029) for financial support to B.B. The AFM data were recorded at the AFM facility of the London Centre for Nanotechnology, with financial support by EPSRC (EP/M028100/1). We acknowledge Richard Thorogate for technical support and members of the Hoogenboom lab for advice and discussions. We thank Rina Rosenzweig at the Weizmann Institute for providing ΔJ-DNAJB1.

## Author contributions

JB, JM, AW, ECJ performed experiments and analysed data. HRS, BB, AW, BWH, JB, JM, AJR designed experiments. All authors contributed to writing the manuscript.

## Notes

### Competing Interest Statement

The authors have declared no competing interest.

