## Supplementary tables, figures and video legends for "Cooperative Amyloid Fibre Binding and Disassembly by the Hsp70 disaggregase"

| Fibre number | Starting length (nm) | Final length (nm) |
|---|---|---|
| 1 | 262 | 177 |
| 2 | 310 | 0 |
| 3 | 888 | 352 |
| 4 | 710 | 360 |
| 5 | 1200 | 968 |
| 6 | 237 | 0 |
| 7 | 374 | 57 |
| 8 | 359 | 0 |
| 9 | 415 | 0 |
| 10 | 556 | 0 |
| 11 | 337 | 230 |
| 12 | 417 | 195 |
| 13 | 152 | 0 |
| 14 | 231 | 0 |
| 15 | 173 | 0 |
| 16 | 1800 | 1380 |
| 17 | 1210 | 0 |

**Supplementary table 1: The starting and final lengths of all αSyn fibres whose disassembly was visualised in AFM videos.**

| Buffer | Buffer Composition |
|---|---|
| HKMD buffer | 50 mM HEPES, 50 mM KCl, 5 mM $MgCl_2$, 2 mM DTT, pH 7.50 |
| HKMT buffer | 50 mM HEPES, 50 mM KCl, 5 mM $MgCl_2$, 2 mM TCEP, pH 7.50 |
| Disaggregation buffer | 50 mM HEPES, 50 mM KCl, 5 mM $MgCl_2$, 2 mM DTT, pH 7.50 5 mM ATP, 6 mM PEP, 20 ng/µL pyruvate kinase |
| Hsp70/110 lysis buffer | 50 mM HEPES, 150 mM KCl, 5 mM $MgCl_2$, 2 mM DTT, EDTA free protease inhibitor tablet (Roche), pH 7.50 |
| Hsp70/110 wash 1 buffer | 50 mM HEPES, 150 mM KCl, 5 mM $MgCl_2$, 0.1 mM PMSF, 2 mM DTT, pH 7.50 |
| Hsp70/110 wash 2 buffer | 50 mM HEPES, 150 mM KCl, 5 mM $MgCl_2$, 40 mM imidazole, 2 mM DTT, pH 7.50 |
| ATP wash buffer | 50 mM HEPES, 150 mM KCl, 5 mM $MgCl_2$, 5mM ATP, 2 mM DTT, pH 7.50 |
| DNAJ lysis buffer | 50 mM HEPES, 750 mM KCl, 5 mM $MgCl_2$, 10 % glycerol, pH 7.50 |
| DNAJ wash 1 buffer | 50 mM HEPES, 150 mM KCl, 5 mM $MgCl_2$, 10 % glycerol, 40 mM Imidazole, pH 7.50 |
| DNAJ wash 2 buffer | 50 mM HEPES, 50 mM KCl, 5 mM $MgCl_2$, 10 % glycerol, 40 mM, Imidazole, pH 7.50 |
| DNAJ elution buffer | 50 mM HEPES, 750 mM KCl, 5 mM $MgCl_2$, 10 % glycerol, 500 mM Imidazole, pH 7.50 |
| DNAJ SEC buffer | 50 mM HEPES, 750 mM KCl, 5 mM $MgCl_2$, 10 % glycerol, pH 7.50 |
| αSyn lysis buffer | 100 mM Tris-HCl, 10 mM EDTA, 2 mM DTT, pH 8.0 |
| αSyn buffer A | 25 mM Tris-HCl, 2 mM DTT, pH 7.7 |
| αSyn buffer B | 25 mM Tris-HCl, 1 M NaCl, 2 mM DTT, pH 7.7 |
| αSyn fibrillation buffer | 50 mM $NaPO_4$, 100 mM NaCl, 0.05% w/v $NaN_3$, pH 7.30 |
| TBS Buffer | 50 mM Tris-HCl, 150 mM NaCl, pH 7.50 |
| HEPES Buffer | 50 mM HEPES, 2mM DTT, pH 7.50 |

**Supplementary table 2. The buffers used for protein purification and amyloid fibre disaggregation reactions**.

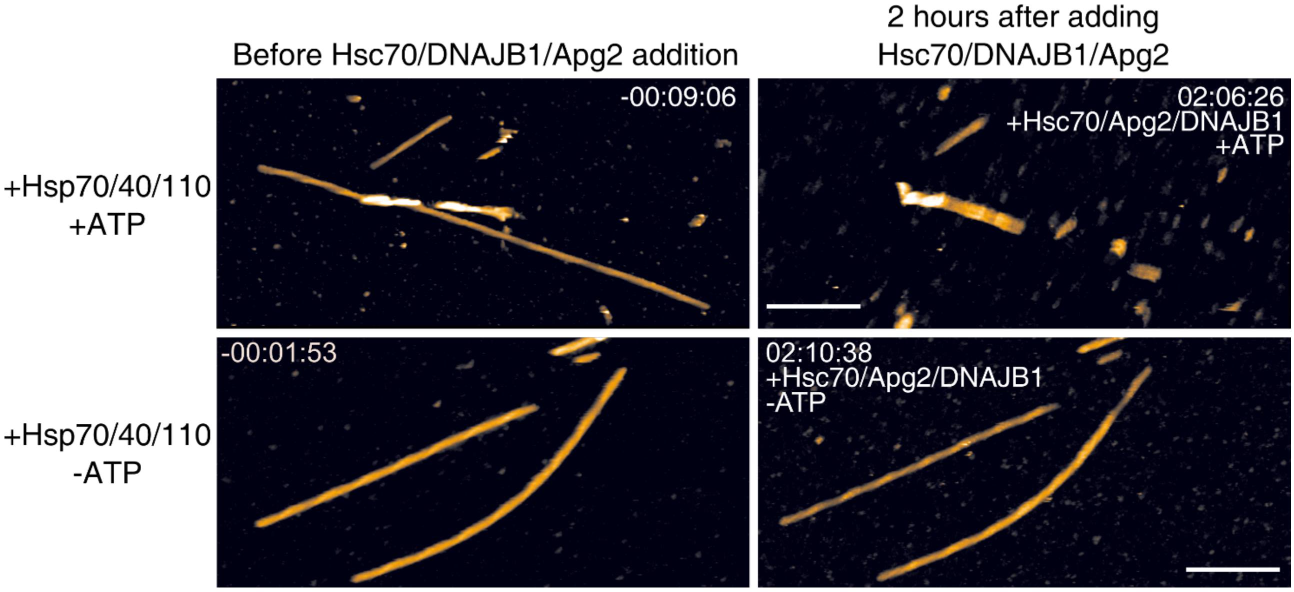


**Supplementary Figure 1. Disaggregation seen in AFM videos requires ATP**. Snapshots of an AFM time-series showing αSyn fibres before addition of chaperones (left panels) and after incubation with chaperones for 2 hours (right panels) in the presence (top panels) and absence (bottom panels) of ATP. The scale bars represent 250 nm.

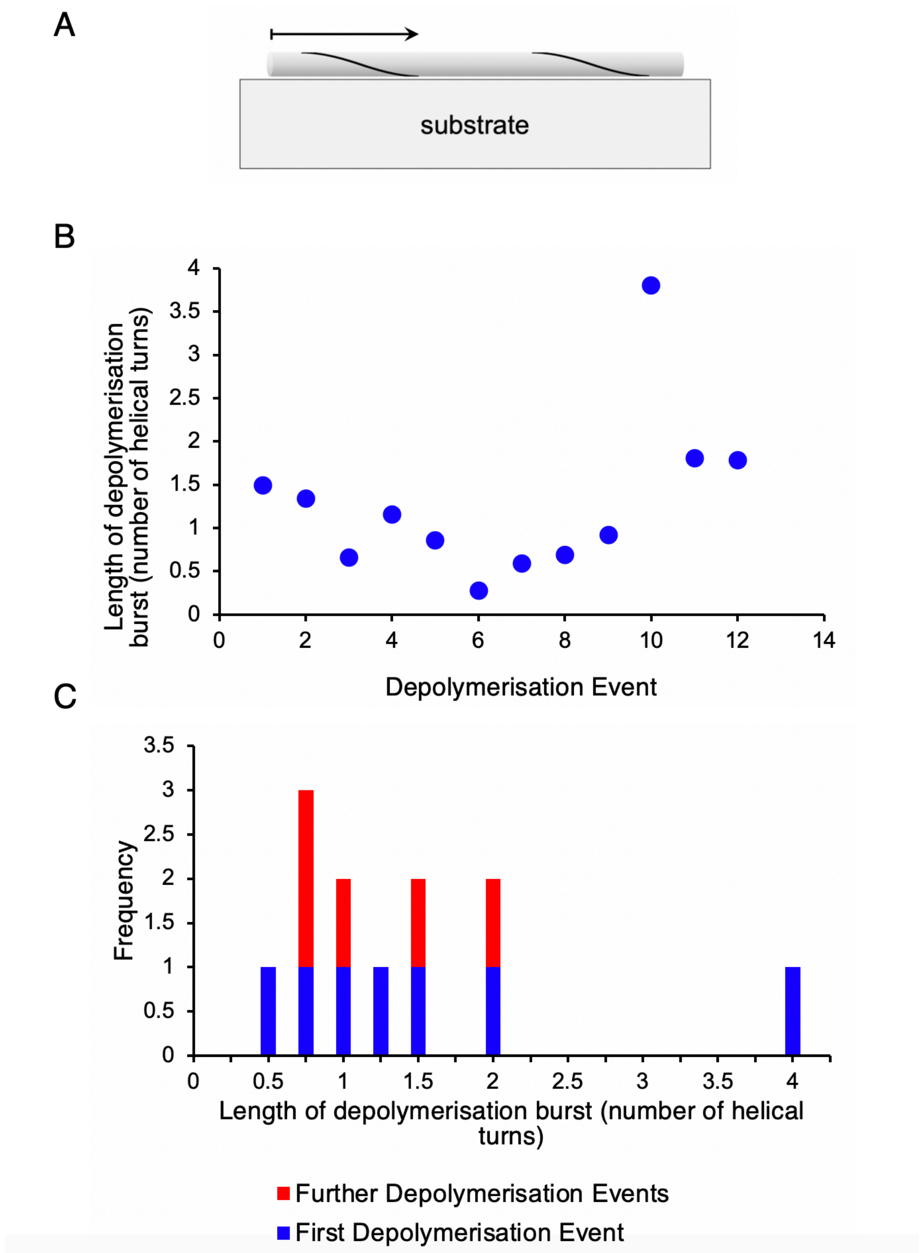


**Supplementary Figure 2. The helical periodicity does not account for the bursts of disassembly**. The chaperone binding sites are on the flexible αSyn termini, which are expected to follow the helical path of the fibre structure (A). We examined whether local attachment to the substrate might arrest disaggregation and account for the burst-like depolymerisation events observed by AFM. In this case, we would expect to find an effect of the helical repeat, visible as a height alternation in the AFM images (Figure 2B), on the positions where the disaggregation comes to a halt. With an effect of periodic attachment, we would expect the disaggregated segment to be limited to lengths between 0 and one helical repeat, since the end of the attached fibre can be assumed to be at a random distance within one helical repeat of the point of the helix that is most firmly attached to the substrate. No such relationship is evident between the helical repeat and the positions where disaggregation is arrested (B). A specific prediction in this scenario would be that the first disaggregated segment would be on average ~50% of the length of subsequently disaggregated segments. Again, this is not consistent with the experimental data (C). The lack of relationship implies that the runs of disassembly are not terminated by periodically less accessible sites on the fibres.

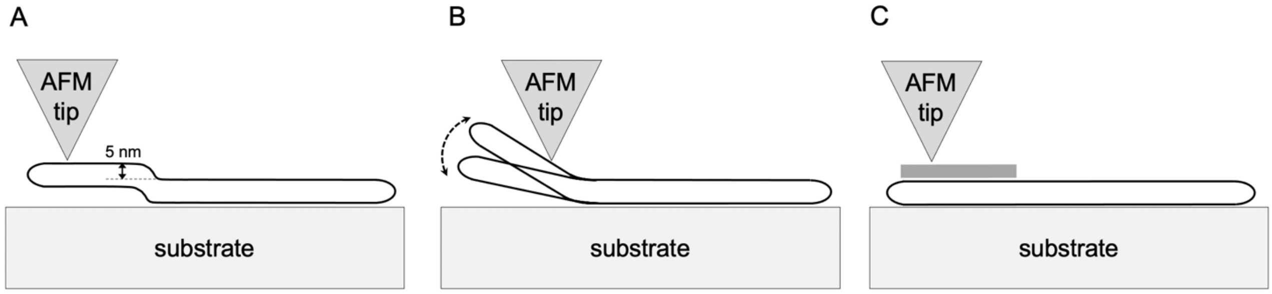


**Supplementary Figure 3. Alternative explanations of the fibre height increase seen by AFM**. The small and uniform height increase preceding a depolymerisation burst does not resemble local detachment from the mica. Local detachment from the mica would not result in a uniform height increase as in A, but would cause increasing mobility of the segment with distance from the detachment point, and a corresponding loss of imaging resolution (B). Dense chaperone binding over the elevated region (C) remains the only plausible explanation accounting for the AFM observations.

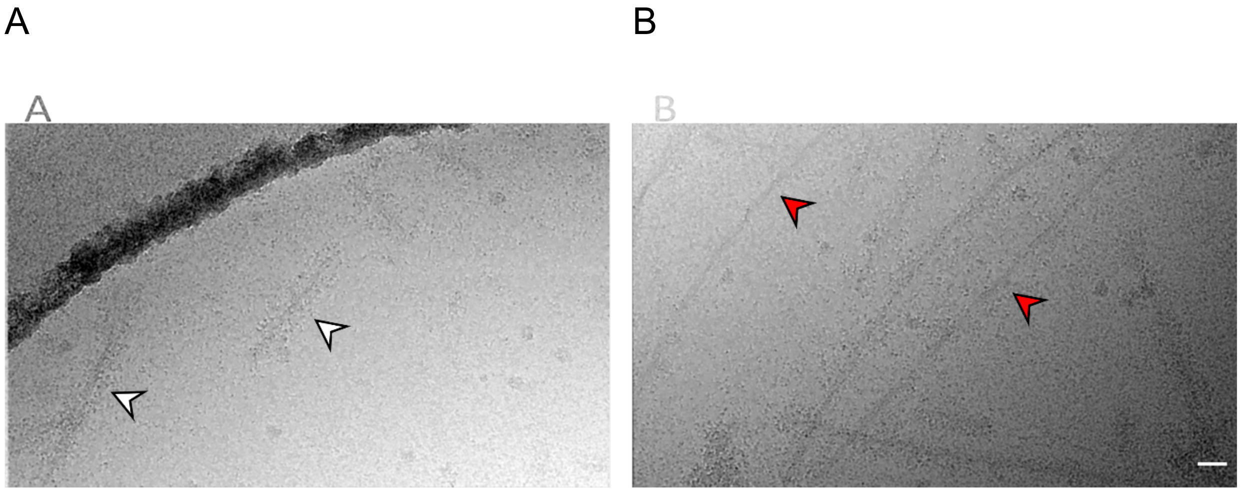


**Supplementary Figure 4. Cryo-EM images of αSyn fibres with DNAJB1, Hsc70, Apg2 and ATP**. A, example of dense binding of chaperones (white arrows), the prominent dark curve at the top left is the edge of the carbon film. B, example of sparser binding (red arrows). The images were denoised to facilitate counting of dense and sparse binding to fibres on a large data set of images from fibres incubated either with the full system or with the system lacking Apg2. Scale bar, 30 nm.

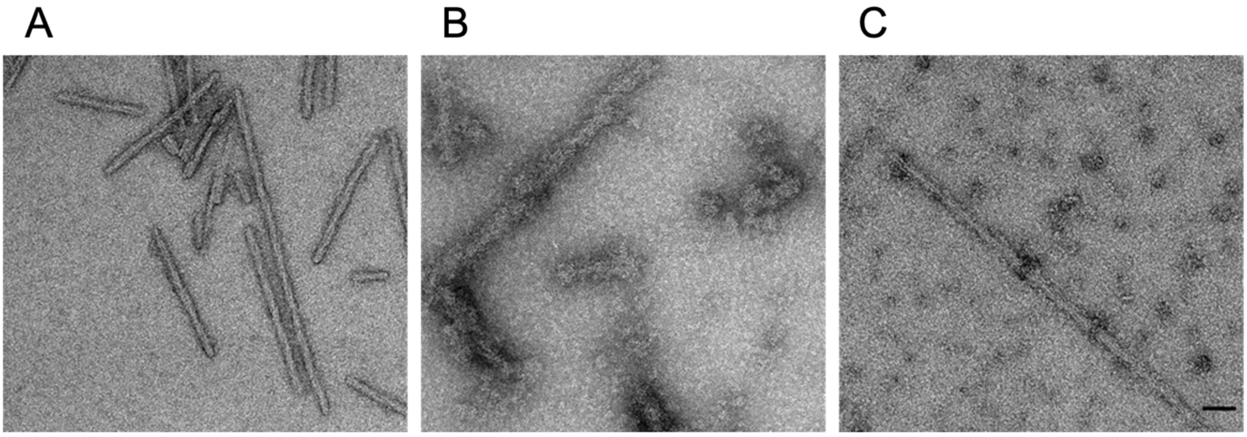

**Supplementary Figure 5. Negative stain EM images showing that chaperone recruitment to fibres did not occur when fibres were incubated with Hsc70/ΔJ-DNAJB1/Apg2/ATP.** In negative stain images, chaperone binding appeared as a pronounced increase in fibre thickness, visible when comparing images of αSyn samples alone to those incubated with Hsc70/DNAJB1/Apg2. A, αSyn fibres alone; B, fibres + Hsc70/DNAJB1/Apg2/ATP; C, fibres + Hsc70/ΔJ-DNAJB1/Apg2/ATP. Scale bar, 50 nm.

**Supplementary videos**

Video 1. AFM movie from Figure 1A.

Video 2. AFM movie from Figure 1B.

Video 3. Tomogram of αSyn fibres with DNAJB1, Hsc70 and ATP.

Video 4. Tomogram of αSyn fibres with DNAJB1, Hsc70, Apg2 and ATP.
